# Are reaching and grasping effector-independent? Similarities and differences in reaching and grasping kinematics between the hand and foot

**DOI:** 10.1101/2021.10.26.464901

**Authors:** Yuqi Liu, James Caracoglia, Sriparna Sen, Erez Freud, Ella Striem-Amit

**Affiliations:** Department of Neuroscience, Georgetown University Medical Center, Washington, DC 20057, USA; Institute of Neuroscience, Key Laboratory of Primate Neurobiology, CAS Center for Excellence in Brain Sciences and Intelligence Technology, Chinese Academy of Sciences, Shanghai, China; Division of Graduate Medical Sciences, Boston University Medical Center, Boston, MA 02215, USA; Department of Psychology, York University, Toronto, Ontario M3J 1P3, Canada; Centre for Vision Research, York University, Toronto, Ontario M3J 1P3, Canada

**Keywords:** Grasping, Reaching, Effector-independent, Visuomotor, Motor cortex

## Abstract

While reaching and grasping are highly prevalent *manual* actions, neuroimaging studies provide evidence that their neural representations may be shared between different body parts, i.e. effectors. If these actions are guided by effector-independent mechanisms, similar kinematics should be observed when the action is performed by the hand or by a cortically remote and less experienced effector, such as the foot. We tested this hypothesis with two characteristic components of action: the initial ballistic stage of reaching, and the preshaping of the digits during grasping based on object size. We examined if these kinematic features reflect effector-independent mechanisms by asking participants to reach toward and to grasp objects of different widths with their hand and foot. First, during both reaching and grasping, the velocity profile up to peak velocity matched between the hand and the foot, indicating a shared ballistic acceleration phase. Secondly, maximum grip aperture and time of maximum grip aperture of grasping increased with object size for both effectors, indicating encoding of object size during transport. Differences between the hand and foot were found in the deceleration phase and time of maximum grip aperture, likely due to biomechanical differences and the participants’ inexperience with foot actions. These findings provide evidence for effector-independent visuomotor mechanisms of reaching and grasping that generalize across body parts.

## Introduction

A central question in motor control regards the functional organization of motor cortex. Successfully performing actions requires precise control and coordination of each effector (acting body part), i.e., muscle and joint movements of a body part. Consistently, primary motor cortex contains a large-scale, albeit imprecise, somatotopic organization in which each cortical area selectively controls movements of a given body part (Meier, Aflalo, Kastner, & Graziano, 2008; Penfield & Boldrey, 1937; Zeharia, Hertz, Flash, & Amedi, 2015; Huntley & Jones, 1991; Asanuma & Rosen, 1972; Strick & Preston, 1978). Furthermore, kinematic and muscle synergies (i.e. patterns that reflect covariations among kinematics and muscle activity) during hand movements were found to be encoded by population neural response in primary motor cortex (Gallego, Perich, Miller, & Solla, 2017; Gallego et al., 2018; Leo et al., 2016; Overduin, D’Avella, Roh, Carmena, & Bizzi, 2015). These findings indicate an organization principle that is specific to effector systems.

Beyond the level of effector-specific representations, though, there is evidence for more abstract motor representations that generalize across different body parts. In both primates and humans, separable neural pathways were found between reaching and grasping actions with the hand, with reaching engaging dorsomedial frontoparietal areas and grasping involving ventrolateral frontoparietal areas (Connolly, Andersen, & Goodale, 2003; Culham, Cavina-Pratesi, & Singhal, 2006; Kaas, Stepniewska, & Gharbawie, 2012; Konen, Mruczek, Montoya, & Kastner, 2013; Yttri, Wang, Liu, & Snyder, 2014). Importantly, these pathways are not merely sub-specializations for a somatotopic hand area, as action representations in these frontoparietal areas may not be specific to hand actions. For example, during motor planning (Gallivan, McLean, Smith, & Culham, 2011) or execution (Magri, Fabbri, Caramazza, & Lingnau, 2019), brain areas including premotor cortex and superior parietal lobule encode target location whether individuals reached with their hand or made saccades with the eyes toward it (also see Heed, Beurze, Toni, Röder, & Medendorp, 2011; Heed, Leone, Toni, & Medendorp, 2016; Leoné, Heed, Toni, & Pieter Medendorp, 2014). Common activation or activity pattern during reaching and grasping actions is also found between the two hands (Gallivan, McLean, Flanagan, & Culham, 2013; Haar, Dinstein, Shelef, & Donchin, 2017; Turella, Rumiati, & Lingnau, 2020), between the hand and the mouth (Castiello et al., 2000), between the hand and tools (Umiltà et al., 2008; Gallivan et al., 2013), or between the hand and foot (Heed et al., 2011, 2016; Leoné et al., 2014; Liu, Vannuscorps, Caramazza, & Striem-Amit, 2020). Neurons responding to both hands or hand and eye were also recorded in posterior parietal cortex in non-human primates (Chang, Dickinson, & Snyder., 2008; Diomedi et al., 2020). These findings indicate that some secondary motor areas are organized based on more abstract motor information that are independent of specific effectors.

Importantly, while evidence for effector-independent motor representations in the brain is accumulating, it is less clear how these are translated into action behavior. If the brain represents motor programs at abstract levels that generalize across muscle groups and body parts (“motor engram”, Lashley, 1950), common motor patterns should be observed across different body parts (“motor equivalence”, Stelmach & Diggles, 1982; Merton, 1972). To examine what information may be represented by effector-independent neural substrates, past studies tested whether motor kinematics features are similar between effectors, specifically if hand-like kinematics extend to other body parts. The kinematic profile and features of hand-reaching and grasping actions have been well characterized: Hand-grasping action is commonly characterized by a transport component where the hand is directed to the target object, and a prehension component by which the hand grasps the object (Jeannerod, 1984). By analyzing tangential hand velocity over time, studies have shown that the transport component typically consists of first, an acceleration phase and then, a deceleration phase as the hand approaches the object (Jeannerod 1984, 1986). The deceleration typically increases with accuracy demands (e.g. smaller objects) whereas the acceleration remains invariant (Gentilucci et al., 1991; MacKenzie et al., 1987; Maitra, Philips, & Rice, 2010; Marteniuk et al., 1990; Jeannerod 1984, 1986). Given these findings, past models propose that the acceleration phase reflects a ballistic movement guided by motor planning prior to movement onset based on the spatial location of the target, whereas the later deceleration phase is more strongly affected by online feedback control (MacKenzie, Marteniuk, Dugas, Liske, & Eickmeier, 1987; Marteniuk, Leavitt, MacKenzie, & Athenes, 1990; Woodworth, 1899; Arbib 1981). Many of these properties extend between the two hands as well as to hand-held tools: The velocity profile, especially during time up to peak deceleration, matched between grasping with either hand and with two hands (Tresilian & Stelmach, 1997; Grosskopf & Khutz-Buschbeck, 2006; Nelson, Berthier, & Konidaris, 2006) and between grasping with the hand or a tool (Gentilucci, Roy, & Stefanini, 2004), suggesting an effector-independent ballistic component prior to the final feedback-control phase (MacKenzie et al., 1987; Marteniuk et al., 1990). In addition, percentage deceleration time increased with smaller (i.e. harder to grasp) objects for both grasping with the hand and with a tool (Gentilucci et al., 2004), suggesting a common mechanism by which object size influences the online feedback control stage.

In terms of prehension, during hand transport, the fingers first extend to a maximum grip aperture that is wider than the object, then close as the hand reaches near to the target and grasp (Jeannerod, 1984, 1986). Importantly, maximum grip aperture robustly scales with object size (Goodale, 1991; Chieffi & Gentilucci, 1993; Freud, Ganel, Avidan, & Gilaie-Dotan, 2016; Freud & Ganel, 2015; Westwood, Danckert, Servos, & Goodale, 2002), indicating encoding of object property (i.e. size) already during the transport stage. This feature has also been established regardless of grasping with one hand or with two hands (Tresilian & Stelmach, 1997) or whether grasping with the hand or a tool (Gentilucci et al., 2004; Itaguchi & Fukuzawa, 2014; but see Maitra et al., 2010), although differed when grasping (to chew) with the mouth (Quinlan & Culham, 2015).

Despite supportive evidence for motor control extending beyond the hand, the origin and generalizability of effector-independence remains unclear. While studies found shared kinematic features between the two hands and between unimanual and bimanual grasping (Grosskopf & Kuhtz-Buschbeck, 2006; Nelson et al., 2006; Tresilian & Stelmach, 1997), it is unclear if this effector-independence is an apriori trait of actions or the extension of a representation originally specific to one hand to the other hand via cross-hemispheric connections that mediate bimanual coordination (Brus-Ramer, Carmel, & Martin, 2009; de Oliveira et al., 2001; Luizzi et al., 2011). Similarly, hand-based coordination experience could result in generalization of originally hand-specific action representation to hand-held tools, eye and mouth.

If action information is represented at an effector-independent level regardless of hand-based coordination, similar kinematics should be observed between the hand and a cortically distant and inexperienced effector. To test this hypothesis, we asked participants to perform reaching and grasping actions with either the hand or the foot. Although the foot has the potential to grasp a object with the big toe and second toe, as used by non-human primates, participants had almost no experience with this action. Testing the foot hence allows us to address whether common kinematic features develop as generalization derived from joint experience between the hand and additional body part (i.e. hand-based interaction) or the existance of an effector-independent motor representation that is generalizable even in an unused effector. We expect such motor representations to manifest in effector-independent kinematics in the initial ballistic stages of movement, and the capacity to scale the aperture based on object size. Importantly, given the participants’ inexperience with foot grasping, we anticipated possible effects of difficulty, such as longer deceleration time (Gentilucci et al., 2004). Further, the hand and foot have different anatomical structures, which affect their movement capacity. The hand can perform opposition between the thumb and other fingers, allowing for precision grasping, whereas the foot is limited (Rolian et al., 2009). In addition, the fingers are more flexible and can move individually whereas the toes cannot (Dempsey-Jones, Wesselink, Friedman, & Makin, 2019). Relately, the foot is biomechanically more constrained in how much separation can be made between the big toe and the second toe, whereas the fingers can separate by a larger amount. Given differences in biomechanical structure and experience, we also expected to observe different kinematics between hand and foot, particularly for action parameters affected by task difficulty.

Specifically, the foot may show a longer deceleration phase driven by difficulty in online motor control (Gentilucci et al., 2004), and may form an overall smaller aperture size during grasping, given the shorter digits. However, similarities in initial ballistic stages of movement and the qualitative capacity of scaling maximum aperture size with object size, despite biomechanical and experiential differences, would provide evidence for effector-independent motor control mechanisms.

## Methods

### Participants

Fifteen participants (3 male, mean age: 29.8 years, SD = 8.9 years) participated in this study. One participant was excluded due to technical issues with the motion capture system, leading to fourteen participants in the reaching task. Two additional participants were excluded from participation in the grasping task because of an inability to separate the big toe and adjacent toe of the tested (right) foot from its resting position, resulting in twelve participants in the grasping task. All participants were neurotypical, self-reported right-handed adults with normal or corrected-to-normal vision and no history of impaired mobility or neurological disorder. Although we did not assess handedness using questionnaire-based tools, self-categorization shows high consistency with questionnaire-based evaluation, especially in right-handed individuals (Chapman & Chapman, 1987). Participants provided written informed consent prior to participation and received a monetary reimbursement. The experimental protocol was approved by the Georgetown University Medical Center Institutional Review Board.

### Apparatus and Stimuli

Participants sat in front of a table on which a target object was positioned at a viewing distance of 30cm. To ensure that the hand and foot can comfortably lay prone on the table, participants sat on a lower chair for use of the hand and sat atop a separate table for use of the foot. Target objects were Efron blocks (Efron, 1969) of different widths (small: 5mm, medium: 10mm, large: 15mm) and heights while matched for surface area (25cm^2^), depth, mass, texture and color.

Movement of the hand and the foot was tracked using an Optitrak 13W motion capture 6-camera system (NaturalPoint, OR, USA). One infrared light-emitting diode (LED) was attached to the side near the tip of the first and second digit of the participant’s right hand and right foot (**Figure 1**). This allowed tracking the change in aperture of the fingers without interfering with natural movement. The motion capture system tracked the three-dimensional (3D) position of each LED at a sampling rate of 100Hz and with a positional accuracy of 0.3mm. The LEDs were arranged in a manner that permitted uninterrupted movement of both the fingers and toes.

**Figure 1.**
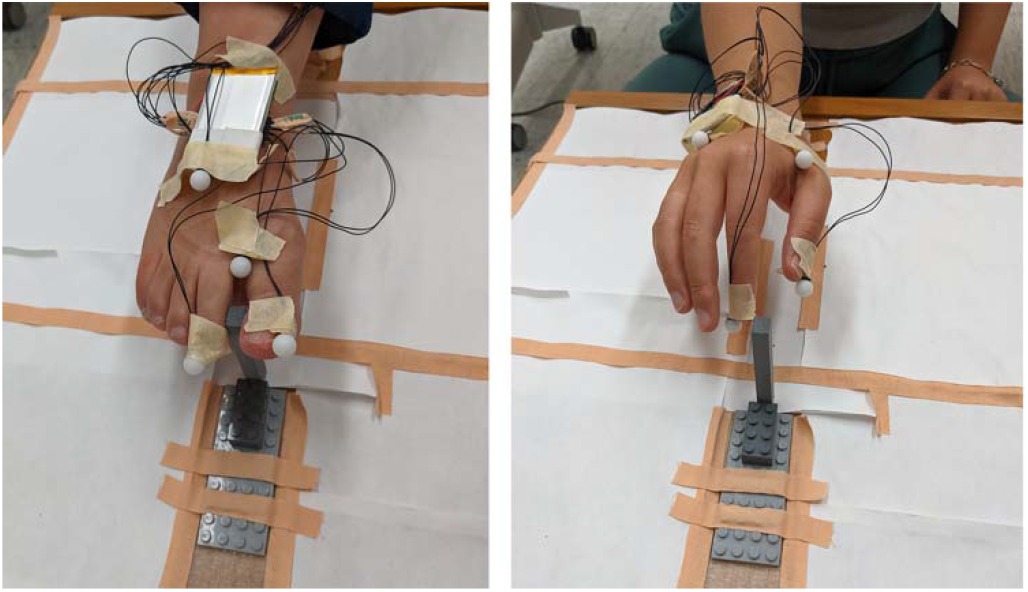
Setup of the sensors and stimuli. For both the hand and foot, one LED sensor was attached to the tip of each of the first two digits. Efron blocks were used as objects.

### Procedure

Each participant performed a reaching and a grasping task in successive order that was counterbalanced across participants. Each task began with six practice trials with each effector (right hand and right foot).

Reaching: At the beginning of each trial, the participant rested the thumb and the index finger (or the big toe and the second toe on the foot) at home position defined by a cube block with the digits touching each other. Upon an auditory “Go” cue, they were instructed to reach toward and touch the front top edge of the target object with the tip of the fingers/toes at a normal pace and then return to the start block. Only the large object (15mm width) was used. The object remained at the target position throughout the task. Each effector was tested in one block of 30 trials. The effector (hand and foot) used in the first block of each task was counterbalanced across participants.

Grasping: The procedure is similar to the reaching task except that upon the “Go” cue, the participant reached toward and grasped the width of the target object using either the thumb and index finger on the right hand, or the big toe and second toe on the right foot (digits 1 and 2 in both cases; **Figure 1**), as if they intended to pick it up, without actually lifting it. Each effector was tested in two blocks in an ABBA design. An object of each width was tested in 15 trials in randomized order within each block. An experimenter switched the target object after each trial and only the target object was visible to the participant in each trial. Since the fingers and toes may differ in the ability of forming aperture, which in turn could affect grasping kinematics, we additionally measured the maximum possible aperture size by asking the participants to extend their index finger and thumb, and the big toe and second toe, to the maximum possible amount. Maximum possible aperture size from two participants was missing due to technical errors.

### Data analysis

3D trajectory data from the motion capture system were preprocessed to extract kinematic information. Average location between the LEDs on the two digits was calculated as a proxy of hand position, based on which we calculated hand and foot velocity. For each trial, movement onset was determined as the point in time when the velocity reached 5% maximum velocity of that trial for at least five consecutive frames (50ms; as done in Schettino, Adamovich, & Poizner, 2003; Ambron, Schettino, Coyle, Jax, & Coslett, 2017; Ganel, Freud, Chajut, & Algom, 2012).

Since actions were performed by first lifting the hand/foot and then landing on the object, movement offset was identified as the point when the fingers/toes were at the furthest position from the body and the lowest position around the end of the movement (peak y and minimum z values respectively). Trials were excluded if have missing data points (i.e. LED sensors not captured by the cameras) or if velocity trajectories indicated unsuccessful grasps, characterized by more than one approaches (y-axis peak) towards the object and re-try at grasping the object itself (frequent also in the absence of visual feedback; Karl, Sacrey, Doan, & Whishaw, 2012). Specific kinematic parameters analyzed for each task are described below:

Reaching: As in past literature, we measured movement duration from the onset to the offset, peak velocity, and absolute time to peak velocity, i.e. the length of the acceleration phase (Jeannerod, 1984; Jeannerod, 1986; Gentilucci et al., 1991; Quinlan & Culham, 2015; Tresilian & Stelmach, 1997). Given evidence that task difficulty can specifically influence time after peak deceleration (Gentilucci et al., 1991; MacKenzie et al., 1987; Marteniuk et al., 1990), we further broke the deceleration phase into absolute time from peak velocity to peak deceleration and absolute time after peak deceleration, as well as measured peak deceleration. Finally, past studies evaluated whether different velocity profiles belong to the same scalar family of curve, hence have identical shape, by testing if the relative proportion of acceleration and deceleration phase remains constant (Gentilucci et al., 1991; MacKenzie et al., 1987). We therefore calculated percentage time to peak velocity relative to movement duration to evaluate the overall shape of velocity profile.

All statistical analyses were performed in JASP (Version 0.10.2) and SPSS (IBM SPSS Statistics for Windows, Version 27.0). For each dependent variable, a paired t-test analysis was performed between hand and foot. Because dependent variables were chosen to test hypotheses regarding specific mechanisms of actions, no multiple-comparisons corrections were performed (Perneger, 1998). Since supporting the effector-independence hypothesis relies on null effects, this approach is more conservative in inferring the effector-independent kinematic properties. Post-hoc analyses were Bonferroni corrected. Moreover, on critical null effects that indicate effector-independence, we performed Bayesian statistical tests and reported BF_10_, i.e. the likelihood of the alternative hypothesis relative to the null hypothesis (Jeffreys 1998, Rouder, Speckman et al. 2009). BF_10_ of less than one indicates the alternative hypothesis is no more likely than the null hypothesis, whereas BF_10_ of less than 1/3 provides support to the null hypothesis over the alternative (Rouder et al., 2009).

Grasping: A 2 (effector: hand, foot) by 3 (object size) repeated-measured ANOVA was performed on each dependent variable. First, we analyzed the same dependent variables as in the reaching task to examine the transport component. In cases where we found an effect of effector in reaching but not grasping or vice versa, we performed a 2 (task: reaching, grasping) by 2 (effector: hand, foot) repeated-measures ANOVA to test the inconsistency between tasks, focusing on the large (15mm) object that was used in both tasks. A significant task × effector interaction effect would reveal differential effects of effector across tasks, indicating influences of the prehension component on transport.

We then examined the prehension component by analyzing maximum grip aperture (MGA) and absolute time to MGA (Jeannerod 1984, 1986; Freud & Ganel, 2015; Ganel et al., 2012). It is possible that some conditions result in both longer movement duration and longer absolute time to MGA (e.g., for larger objects, Gentilucci et al., 1991). We therefore also analyzed percentage time to MGA to compare the temporal structure of the manipulation component. Additionally, to measure sensitivity to object size, past studies calculated the linear slope between aperture size and object size, with a slope larger than zero signaling sensitive scaling (Goodale, 1991; Jeannerod, 1986; Freud et al., 2016). Thus, in addition to running an ANOVA on the MGA, we tested whether the slope between MGA and object size is larger than zero for each effector.

Finally, to determine how early sensitivity to object size arises in the hand and foot, we calculated the linear slope between aperture size and object size at each frame after resampling all trials to the same length (see below), and at 11 normalized movement time points (Freud et al., 2016; from 0% to 100% in 10% steps).

For visualization purposes, we resampled the velocity and aperture profile from each trial to the group average movement duration of each condition, preserving the shape and magnitude of the original time course (Quinlan & Culham, 2009). Then we averaged all trials within each condition to show group-level velocity and aperture profile in absolute time (in seconds). We additionally sampled the velocity and aperture size at each normalized time point (Freud, Culham, Namdar, & Behrmann, 2019; Freud & Ganel, 2015; Ganel et al., 2012) to plot the velocity profile of the hand and foot at an aligned time scale.

## Results

### Reaching

Overall 4.9% of trials were excluded, with no difference between hand (*M* = 5.0%, *SE* = 2.3%) and foot trials (*M* = 4.8%, *SE* = 2.0%; t(13) = 0.09, *p* = .932).

The velocity profiles of hand and foot reaching movements were remarkably similar, both consisting of an acceleration phase followed by deceleration, with visually similar peak velocity and time to peak velocity (**Figure 2**). **Table 1** summarizes the result of each dependent variable (mean for hand and foot, t-test statistics and Bayes factors). First, movement duration was significantly longer for the foot than the hand (t(13) = 2.27, *p* = .041; see detail in **Table 1.1**). This effect is driven by a statistically significantly longer time after peak deceleration for the foot vs. hand (t(13) = 2.61, *p* = .021; **Table 1.5**), with no difference in absolute time to peak velocity (t(13) = 0.62, *p* = .549, BF_10_ = 0.32; **Table 1.2, Figure 2A**) or peak velocity to peak deceleration (t(13) = 0.77, *p* = .457, BF_10_ = 0.35; **Table 1.4**). Absolute time to peak velocity offers support to the null hypothesis of similarity between the hand and foot. This finding indicates that the foot and hand differ only during the final feedback-controlled stage (MacKenzie et al., 1987; Marteniuk et al., 1990; Woodworth, 1899; Arbib 1981), but not during the earlier ballistic movement stage.

**Table 1.** Summary of the results from the reaching task

| Transport parameter |  | Hand | Foot | t-test | BF<sub>10</sub> |
| --- | --- | --- | --- | --- | --- |
|  |  | (Mean (SE)) | (Mean (SE)) | (t(13), p) |  |
| 1 | Movement duration (s) | 1.05 (0.05) | 1.16 (0.04) | 2.27, .041* |  |
| 2 | Time to peak velocity (s) | 0.310 (0.017) | 0.321 (0.013) | 0.62, .549 | 0.32† |
| 3 | Time to peak velocity (%) | 30.4 (1.68) | 28.0 (1.00) | 1.48, .163 | 0.66 |
| 4 | Peak velocity to peak deceleration (s) | 0.335 (0.054) | 0.289 (0.025) | 0.77, .457 | 0.35 |
| 5 | Time after peak deceleration (s) | 0.403 (0.060) | 0.567 (0.030) | 2.61, .021* |  |
| 6 | Peak deceleration (mm/s<sup>2</sup>) | -9797.4 (1526.9) | -11794.2 (1306.4) | 1.01, .329 | 0.42 |
| 7 | Peak velocity (mm/s) | 771.4 (38.4) | 847.8 (23.1) | 2.05, .062 | 2 |

\*: Significant t-tests supporting difference between hand and foot.
†: BF<sub>10</sub> less than 1/3 supporting no difference between hand and foot.

**Figure 2.**
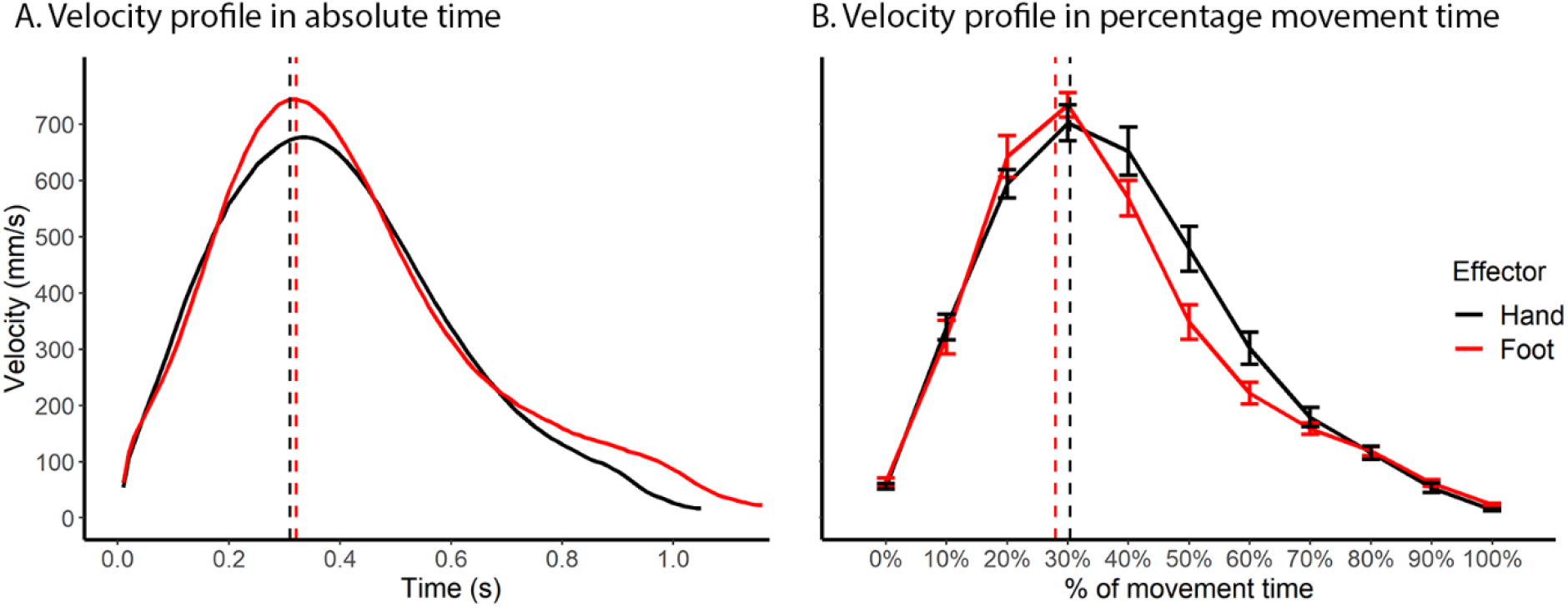
Comparison of the reaching velocity profile between the hand and the foot. **A**. Velocity profile as a function of absolute time. **B**. Velocity profile over percentage movement time. Vertical lines denote time of peak velocity calculated from raw, unnormalized data. Overall the hand and the foot showed remarkably similar velocity profiles, especially from movement onset to peak velocity. Error bars denote standard errors.

Despite difference in absolute movement time, the percentage time to peak velocity of the entire movement duration did not differ between the hand and foot (t(13) = 1.48, *p* = .163, BF_10_=0.66; **Table 1.3, Figure 2B**), suggesting overall consistent reaching movement structure between hand and foot. Finally, there was no difference between the foot and hand in terms of peak deceleration (t(13) = 1.01, *p* = .329, BF_10_ = 0.42; **Table 1.7**), whereas results on peak velocity (t(13) = 2.05, *p* = .062, BF_10_ = 2; **Table 1.6**) were ambiguous. Overall, reaching movement kinematics was similar between the hand and foot, with the early, ballistic component of the action showing consistency, supporting an effector-independent motor control mechanism, whereas differences in time course arising only in the later deceleration stage.

### Grasping

Overall 12.3% of trials was excluded, with more foot trials (*M* = 20.0%, *SE* = 2.4%) excluded than hand trial (*M* = 4.5%, *SE* = 2.5%; t(11) = 5.31, *p* < .001), indicating the difficulty of foot grasping.

#### Transport component

Across all dependent variables tested in the transport component (see Methods), there was no main effect of object size (*p*s > .070) and only one significant interaction between object size and effector on movement duration (see detail below). Therefore, we primarily report the effect of effector on the transport variables, focusing on our main question of whether the foot and hand share similar kinematic properties.

As with the reaching task, both hand and foot showed a bell-shaped velocity profile (**Figure 3**). However, the velocity profile of the foot appears to be more right-skewed, i.e., with a longer deceleration phase, than the hand. Statistical analyses support this observation. First, movement duration was longer for the foot than the hand (F(1,11) = 32.85, *p* < .001; **Table 2.1, Figure 3A**). Subsequent analyses revealed that the time to peak velocity was similar between hand and foot (F(1,11) = 0.08, *p* = .785, BF_10_ = 0.28; **Table 2.2, Figure 3A**), while the foot took a longer time than the hand both from peak velocity to peak deceleration (F(1,11) = 23.70, *p <* .001; **Table 2.4**) and from peak deceleration to the end of the movement (F(1,11) = 24.84, *p* < .001; **Table 2.5**).

**Table 2.** Effect of effector on transport dependent variables in the grasping task

| | **Transport parameter** | Hand<br>(Mean (SE)) | Foot<br>(Mean (SE)) | F-test<br>($F(1,11)$, $p$) | $BF_{10}$ |
| --- | --- | --- | --- | --- | --- |
| 1 | Movement duration (s) | 1.09 (0.05) | 1.58 (0.10) | 32.85, $< .001^*$ | |
| 2 | Time to peak velocity (s) | 0.316 (0.015) | 0.320 (0.018) | 0.08, .785 | 0.28<sup>†</sup> |
| 3 | Time to peak velocity (%) | 29.4 (1.01) | 21.0 (0.93) | 37.77, $< .001^*$ | |
| 4 | Peak velocity to peak deceleration (s) | 0.255 (0.034) | 0.398 (0.020) | 23.70, $< .001^*$ | |
| 5 | Time after peak deceleration (s) | 0.530 (0.037) | 0.955 (0.083) | 24.84, $< .001^*$ | |
| 6 | Peak velocity (mm/s) | 798.6 (36.7) | 777.8 (31.5) | 0.57, .465 | 0.81 |
| 7 | Peak deceleration (mm/s<sup>2</sup>) | -9109.8 (965.0) | -12446.8 (1377.1) | 10.31, .008* |  |

\*: Significant F-tests supporting difference between hand and foot.
<sup>†</sup>: $BF_{10}$ less than 1/3 supporting no difference between hand and foot.

**Figure 3.**
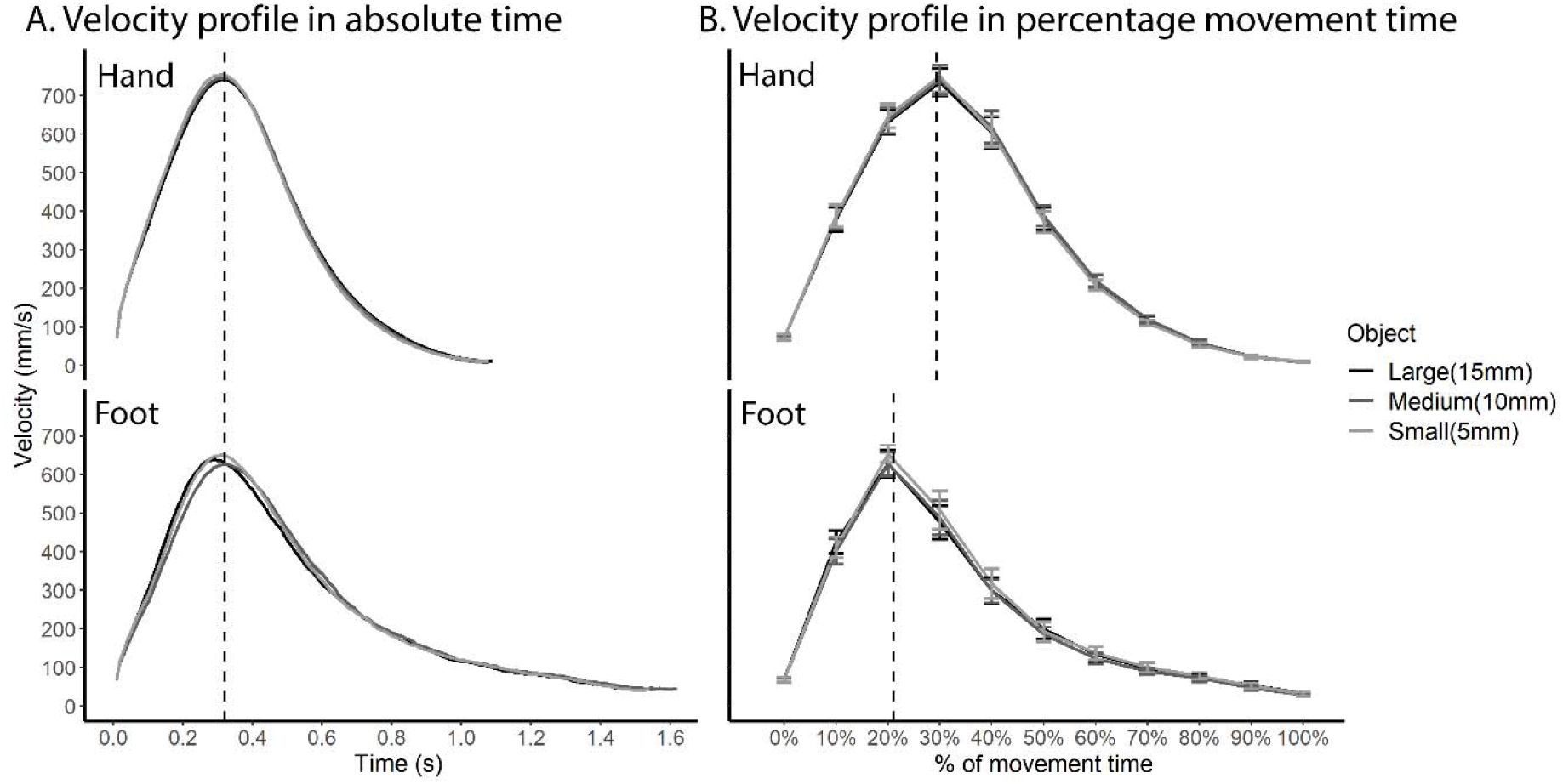
Comparison of the velocity profile between the hand and the foot in the grasping task. **A**. Velocity profile as a function of absolute time. **B**. Velocity profile over percentage movement time. Vertical lines denote time of peak velocity calculated from raw, unnormalized data, averaged across object sizes given no main effect of object size. The hand and the foot showed remarkably similar velocity profiles in absolute time up to peak velocity, with a longer deceleration phase for the foot. Error bars denote standard errors.

As a result of a prolonged deceleration phase for the foot, the proportion of the acceleration phase, i.e. percentage time to peak velocity, was smaller for the foot than the hand (F(1,11) = 37.77, *p* < .001; **Table 2.3, Figure 3B**). Consequently, the shape of the velocity profile of hand and foot was not identical.

As with the reaching task, there was no difference in peak velocity between hand and foot (F(1,11) = 0.57, *p* = .465, BF_10_ = 0.81; **Table 2.2**). However, the foot showed a larger peak deceleration than the hand (F(1,11) = 10.31, *p* = .008; **Table 2.7**). The longer deceleration phase and larger peak deceleration may reflect difficulty in grasping with foot so that the movement had to be performed more slowly.

Finally, there was a significant interaction between effector and object size (F(2,22) = 4.88, p = .018) on movement duration. Whereas object size did not affect movement duration in the hand condition (*p* = .563), the medium object (*M* = 1.62s, *SE* = 10.74s) resulted in a longer movement time than the small object (*M* = 1.54s, *SE* = 10.22s) in the foot condition (*p* < .008, Bonferroni corrected), with no difference between the large object (*M* = 1.56s, *SE* = 11.38s) and the medium or small object.

Overall, as in reaching, the transport of the foot towards the grasped target was again similar to that of the hand for the earlier stages of movement until peak velocity, whereas difference between the hand and foot occurred during the deceleration stage in a manner partly dependent on the intended grasped target size.

For some dependent variables, e.g., percentage time to peak velocity and absolute time from peak velocity to peak deceleration, there was no effect of effector in reaching but an effect in grasping. We performed subsequent post-hoc analyses with task (reaching, grasping) and effector (hand, foot) as independent variables to test these task differences (**Table 3**; Bonferroni corrected for multiple comparisons). Regarding percentage time to peak velocity, there was an interaction between task and effector (F(1,11) = 9.33, *p* = .011), with no difference between hand and foot in reaching (*p* = .146) but a smaller normalized time to peak velocity for foot vs. hand in grasping (*p* < .001). Regarding absolute time from peak velocity to peak deceleration, there was also an interaction between task and effector (F(1,11) = 12.26, *p* = .005), again with no difference between hand and foot in the reaching task (*p* > .999) but a longer time for the foot than the hand in grasping task (*p* = .033). Overall, velocity profile between hand and foot differed more strongly in grasping than reaching.

**Table 3.** Comparing the effect of effector between reaching and grasping

| Transport parameter | | Task Effect<br>($F(1,11), p$) | Reaching<br>(Bonferroni<br>corrected $p$) | Grasping<br>(Bonferroni<br>corrected $p$) |
| --- | --- | --- | --- | --- |
| 1 | Time to peak velocity (%) | 9.33, .011* | .146 | < .001* |
| 2 | Peak velocity to peak<br>deceleration (s) | 12.26, .005* | > .999 | .033* |

\*: Significant F-tests supporting task effect on the effect of effector, and significant post-hoc comparisons between hand and foot.

In summary, while the velocity profile of hand and foot are highly similar in reaching task, the foot showed a more right-skewed velocity profile than the hand in grasping task that was mainly driven by a prolonged deceleration phase after peak velocity. These differences may be caused by an overall greater difficulty in the grasping task for foot compared with hand, or by differential mechanisms between hand and foot in how the transport and prehension components interact.

#### Prehension component

During grasping, the fingers first extend and then close as approaching the object, forming a maximum grip aperture that scales with object size (Chieffi & Gentilucci, 1993; Freud et al., 2016; Freud & Ganel, 2015; Westwood et al., 2002). We analyzed maximum grip aperture (MGA) and its scaling with object size (**Figure 4C, Table 4**). Regarding MGA, the main effect of effector was significant (F(1,11) = 25.07, p < .001; **Table 4.1**), with an overall larger MGA for the hand vs. the foot, as expected given the different digit lengths and dexterity. Indeed, the maximum possible aperture size was significantly larger for hand (*M* = 143.2mm, *SE* = 2.00mm) vs. foot (*M* = 44.8mm, *SE* = 0.98mm; t(9) = 15.00, p < .001), with the foot MGA reaching the maximum aperture limit for all object sizes (**Supplementary figure S1**).

**Table 4.** Effect of effector and object size on prehension dependent variables in the grasping task

| Prehension parameter | | Effect of Effector<br>($F(2,22)$, $p$) | | Effect of Object size<br>($F(2,22)$, $p$) | | | Interaction<br>($F(2,22)$, $p$) |
| --- | --- | --- | --- | --- | --- | --- | --- |
|  |  | Hand<br>(Mean<br>(SE)) | Foot<br>(Mean<br>(SE)) | Large<br>(Mean<br>(SE)) | Medium<br>(Mean<br>(SE)) | Small<br>(Mean<br>(SE)) |  |
| 1 | MGA (mm) | 25.07, $< .001^*$ | | 69.96, $< .001^*$ | | | 26.42, $< .001^*$ |
|  |  | 60.1<br>(2.40) | 43.0<br>(2.49) | 54.4<br>(1.80) | 51.4<br>(1.77) | 48.83<br>(1.77) |  |
| 2 | Time to MGA<br>(s) | 44.29, $< .001^*$ | | 7.54, .003* | | | 2.95, .073 |
|  |  | 0.55<br>(0.068) | 1.15<br>(0.068) | 0.89<br>(0.05) | 0.86<br>(0.05) | 0.81<br>(0.05) |  |
| 3 | Time to MGA<br>(%) | 28.03, $< .001^*$ | | 7.23, .004* | | | 1.10, .352 |
|  |  | 51.6<br>(3.66) | 72.3<br>(1.89) | 63.5<br>(2.34) | 60.7<br>(2.28) | 59.0<br>(2.36) |  |

\*: Significant F-tests

**Figure 4.**
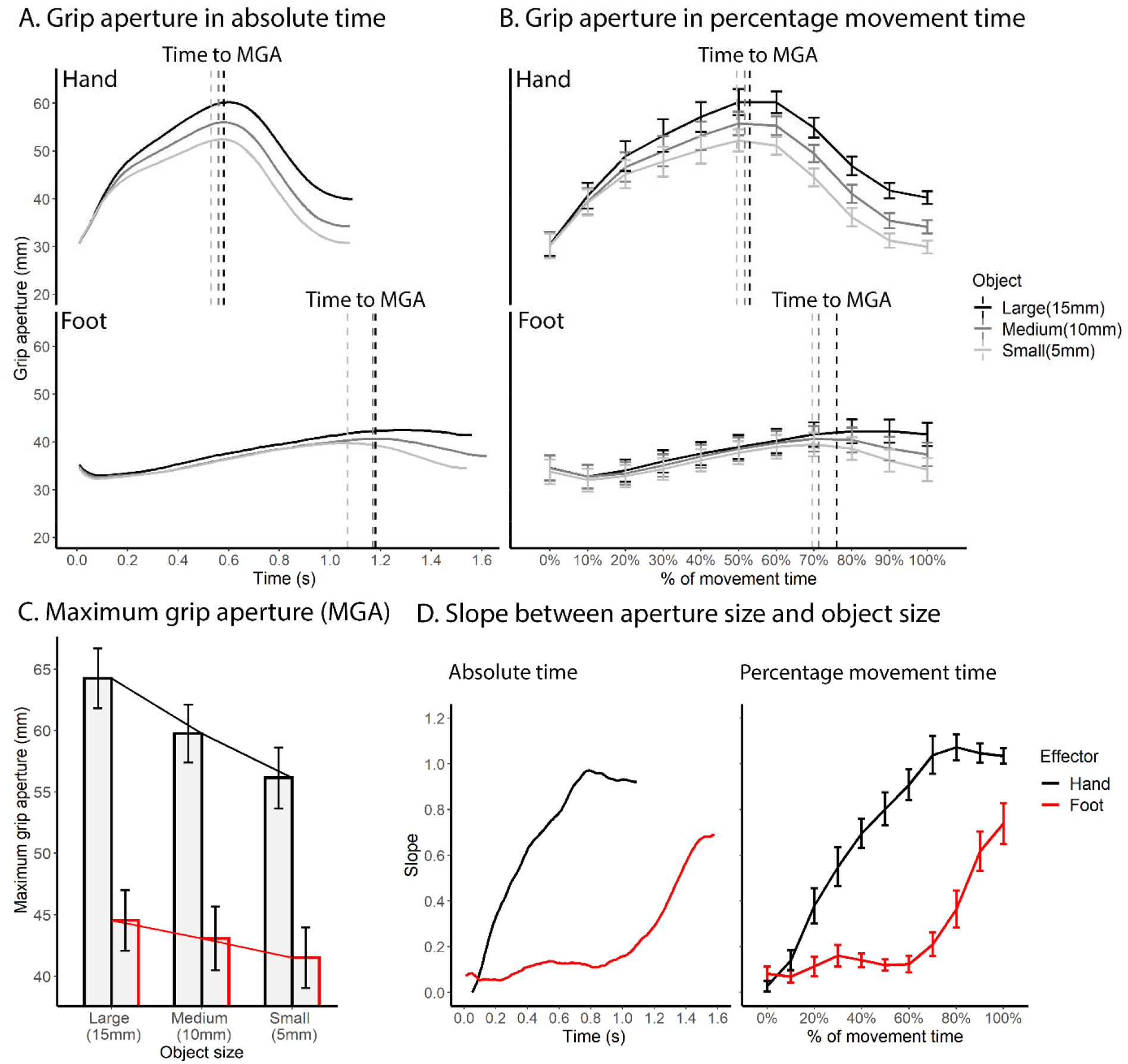
Comparison of the prehension kinematics between the hand and the foot. **A**. Changes in aperture size over absolute time for each effector and object size. The dashed vertical lines denote time to maximum grip aperture. **B**. Changes in aperture size over percentage time for each effector and object size. Vertical dashed lines denote percentage time to maximum grip aperture. **C**. Maximum grip aperture for each effector and object size. A line graph is superimposed on the bar graph to visualize the trend. **D**. Slope between aperture size and object size for each effector over absolute and percentage time. The slope for foot began to rise much later than hand.

Importantly and as expected, despite these anatomical differences, there was a main effect of object size (F(2,22) = 69.96, p < .001; **Table 4.1**). Across both hand and foot, MGA increased with object size (Bonferroni corrected ps < .001 for all post-hoc pair-wise comparisons), consistent with past literature (Chieffi & Gentilucci, 1993; Freud et al., 2016). Finally, there was an interaction between effector and object size (F(2,22) = 26.42, p < .001) driven by a stronger effect of object size in hand vs. foot. This can be seen in subsequent analyses showing significantly above-zero slope between MGA and object size for both the hand and foot (ps < .001, **Figure 4C**), indicating sensitivity and oversizing of both effectors with respect to object size, while the slope was larger for the hand (*M* = 0.81, *SE* = 0.086) vs. the foot (*M* = 0.30, *SE* = 0.067).

We then analyzed time to MGA. As expected based on past research on hand grasping (Jeannerod, 1984; Marteniuk et al., 1990; Tresilian & Stelmach, 1997), MGA occurred after peak velocity for both foot and hand, with later MGA time for larger objects (**Figure 4A** and **4B**). Moreover, MGA occurred later for the foot as compared to the hand. Statistical analyses supported these observations (**Table 4.2**). Regarding absolute time to MGA, there was a significant main effect of effector (F(1,11) = 44.29, *p* < .001), with a later time for foot vs. hand (**Figure 4A**). The main effect of object size was also significant (F(2,22) = 7.54, *p* = .003), with a later MGA for larger objects across effectors, consistent with past findings for hand grasping (Jeannerod, 1984; Marteniuk et al., 1990; Tresilian & Stelmach, 1997). Post-hoc comparisons revealed a significant difference between the large and small object (*p* = .028), with no difference between the medium object and the large or small object (*p*s > .06; see **Table 4.2** for descriptive statistics of each object). There was no interaction between effector and object size (F(2,22) = 2.95, *p* = .073). Analyses of percentage time to MGA led to the same conclusions (**Table 4.3, Figure 4B**), with a later percentage MGA time for the foot vs. hand (F(1,11) = 28.03, p < .001), a main effect of object size driven by a later time for the large vs. small object (F(2,22) = 7.23, p = .004), and no interaction between effector and object size (F(2,22) = 1.10, p = .352).

Finally, to examine the time course of sensitivity, we calculated the linear slope of aperture size to object size over time. As shown in **Figure 4D**, the slope of the aperture began to increase at a much later time point for foot vs. hand in terms of both absolute time and percentage time, indicating that the sensitivity of aperture to object size occurred much later. The same trend can be seen in **Figure 4A** and **4B** where the aperture profile of different object sizes began to show separation early for the hand but rather late for the foot.

Overall, significant differences were found in the timing of sizing sensitivity between the effectors, as well as the differences in aperture size. However, prehension of the foot showed grip aperture properties known to characterize hand prehension, including pre-shaping the digits based on object size and later time of maximum aperture for larger objects.

## Discussion

We examined kinematic properties of hand and foot movement during reaching and grasping actions to investigate what aspects of visually guided actions are shared between effectors and may be controlled by effector-independent motor neural mechanisms. We found similarities in kinematics between foot and hand including *(i)* same absolute time to peak velocity in both reaching (**Table 1.2)** and grasping (**Table 2.3**), along with similar velocity profiles (i.e., the proportion of acceleration and deceleration phase) in the reaching task (**Figure 2**) *(ii)* maximum grip aperture scaled with object size (**Figure 4C**), and *(iii)* later time to maximum grip aperture with larger objects (**Figure 4, Table 4.2**). These findings expand on previous literature showing kinematic consistency across the two hands as well as for hand-held tools, thereby showing that at least some effector-independent kinematic properties can extend to a distant and inexperienced effector. Differences between hand and foot were also found, in *(i)* longer deceleration time for foot vs. hand (**Table 1.5, Table 2.4, 2.5**), *(ii)* later time to maximum grip aperture for foot vs. hand (**Figure 4A, 4B, Table 4.2, 4.3**), *(iii)* smaller scaling of aperture size with object size for foot vs. hand (**Figure 4C**), and *(iv)* later onset of scaling of aperture size with object size for foot vs. hand (**Figure 4D**) These differences could stem from inexperience with foot actions, different biomechanical capacities between the hand and foot, or, alternatively, effector-dependent mechanisms underlying certain properties of these actions. We discuss these findings under the current framework of motor planning and execution.

### Similar ballistic movement phase across hand and foot

It has long been proposed that the earlier phase of reaching and grasping actions are primarily controlled by a motor planning mechanism, whereas the later stages are guided by online feedback control mechanisms (Arbib, 1981; MacKenzie et al., 1987; Marteniuk et al., 1990; Keele, 1968; Dixon & Glover, 2009; Glover, 2002; Glover & Holloway, 2004). Complete reaching movements can be guided by pre-planned motor commands even when later correction is rendered impossible. For instance, in cases where target location is changed suddenly after movement onset yet correction of movement trajectory is hindered by posterior parietal lesion or TMS (Desmurget et al., 1999; Gréa et al., 2002), participants cannot correct the movement trajectory to the new target position. Importantly, however, they still make smooth movements toward the initial target location, indicating pre-planned movement trajectory regardless of online sensory information.

Under typical circumstances, the motor plan seems to be more strongly reflected in the acceleration phase of transport (Elliott et al., 1999; Gentilucci et al., 1991; Mackenzie et al., 1987; Marteniuk et al., 1990). For instance, in hand reaching/grasping studies that introduce unexpected perturbations (e.g., decrease or increase in resistance) following movement initiation, the hand typically requires longer to complete the movement in the perturbation condition(s); however, acceleration kinematic parameters (e.g., peak velocity, absolute time to peak velocity) tend to not differ (Elliott et al., 1999), suggesting that the acceleration phase is less influenced by online sensory information. With respect to our data, we found that in both reaching and grasping, the acceleration phase of transport shares more similarities (i.e., in absolute time to peak velocity) between hand and foot than the deceleration phase, suggesting that the acceleration phase is less dependent on specific sensorimotor parameters associated with each effector. During the acceleration phase of arm/leg movement, both effectors demonstrated similarity in absolute time to peak velocity and in peak velocity. A matched acceleration phase was also reported between grasping with hand or a tool (Gentilucci et al., 2004; but see Maitra et al., 2010). Taken together, our data posits that (ballistic) motor planning for reaching and grasping actions is effector independent, and the motor command may form at a level of abstraction above that of the generation of effector-specific muscle command, i.e., specificity to the hands. At neural level, neuroimaging and single-cell recording studies reported common activation or activity patterns in premotor cortex and posterior parietal cortex during motor planning and execution of reaching/pointing and grasping actions across effectors (Gallivan et al., 2013; Heed et al., 2011, 2016; Leoné et al., 2014; Liu et al., 2020; Diomedi et al., 2020; Magri et al., 2019; Chang et al., 2008; Diomedi et al,. 2020; Gamberini et al., 2011; Ferraina et al., 1997; Hadjidimitrakis et al., 2011). These areas may serve as neural substrates for shared motor control, i.e., the derivation of the shared kinematic properties observed in our study.

### Differences in the deceleration phase between the foot and the hand

Despite similarities in the acceleration phase, a longer deceleration was found for the foot as compared with the hand. In the reaching task, the longer deceleration phase for foot was primarily driven by a longer time from peak deceleration to movement offset. During grasping, the foot took an overall longer absolute time after peak velocity, both from peak velocity to peak deceleration and from peak deceleration to end. The prolonged deceleration phase led to a more right-skewed velocity profile of foot than hand (i.e., shorter percentage time to peak velocity in foot), resulting in different normalized velocity profile shapes. One possibility is that these results reflect differences in neural online control between the effectors, supporting effector-dependent motor control based on somatotopically-selective areas (Yttri et al., 2014), as well as a separate representation type for action planning as compared to its control (Glover 2004). However, it may also stem from increased difficulty and inexperience of foot action. Past studies reported difference in time after peak deceleration for hand movement between different levels of accuracy demands (Marteniuk et al., 1990; Gentilucci et al., 1991; Mackenzie et al., 1990). This final movement stage requires precise coordination between sensory and motor information, heavily relying on online feedback control (Elliott et al., 1999; Gentilucci et al., 1991; Mackenzie et al., 1987; Marteniuk et al., 1990). Therefore, one possibility is that precise motor control of the foot/leg is intrinsically harder due to its heavier weight, more biomechanical constraints and less flexibility in separating the toes for grasping, leading to longer time after peak deceleration.

An alternative but non-exclusive account is that the participants were simply less experienced in grasping objects with the foot, finding this task more difficult. There is evidence that although grasping with uncommon or novel effectors led to longer grasping time and different aperture profile compared with typical grasping with the thumb and index finger, practice can eliminate these differences (Itaguchi, 2020; Bouwsema, van der Sluis, & Bongers, 2014; Itaguchi & Fukuzawa, 2014). Added evidence for the difficulty of foot grasping can be found in the larger number of unsuccessful grasp trials (which were thus excluded from analysis). On this account, foot and hand actions are controlled by common underlying mechanisms, but it requires experience to efficiently translate the motor program to kinematic patterns.

### Similarities and differences in prehension across effectors

The maximum grip aperture scaled with object size for both the foot and the hand. The scaling of maximum grip aperture to object size in grasping has been robustly and widely reported in past studies (Chieffi & Gentilucci, 1993; Jeannerod, 1984a, 1986), and has been measured as indicating sensitivity of grasping motor control to object size (Freud et al., 2016; Westwood et al., 2002). MGA is less affected by visual illusion or perceptual deficits, hence reflects visuomotor functions rather than pure perceptual information (Dixon & Glover, 2009; Freud et al., 2016; Westwood et al., 2002; Aglioti, DeSouza, & Goodale, 1995). As aperture scaling can occur in the absence of visual feedback (Hu, Eagleson, & Goodale, 1999), but typically occurs in later stages of movement, i.e. during deceleration, it is considered to be guided by both online control and planning (Glover, 2004). Moreover, Past studies also found scaling of MGA with object size for unimanual and bimanual grasping (Tresilian & Stelmach, 1997), for grasping with a tool (Gentilucci et al., 2004; but see Maitra et al., 2010), and for grasping/biting with the mouth (Castiello, 1997; Churchill et al., 1999; Quinlan & Culham, 2015), indicating common mechanism by which the grasping motor control system encodes visual information regarding object size prior to contact. We provide additional evidence that such motor control mechanism also applies to foot, despite limited experience in grasping with foot.

Although we found scaling of MGA with object size in both effectors, the foot showed a smaller slope (0.30) between MGA and object size than the hand (0.81; well within previously reported ranges; Freud et al., 2016), indicating lower capacity for scaling the aperture of the foot. Biomechanically, the toes have a more limited movement range than the fingers either due to a shorter length of the toes or less flexible joints, as evident in a larger maximum possible aperture size for the hand as compared with the foot. With these constraints, foot aperture has a ceiling effect to its aperture: its maximal aperture is on average 44.8mm, whereas the MGA was 43.0mm, and may only increase by a limited amount with increased object size (**Supplementary Figure S1**). This therefore manifests in an overall smaller maximum grip aperture for foot vs. hand. Similarly, biomechanical constraints and lack of experience may also have resulted in delayed MGA and later-onset sensitivity to object size (**Figure 4D**) in foot, where the toes could not separate optimally until the speed of the foot reduced to a certain amount. On these possibilities, the mechanism by which object size is encoded during pre-shaping is shared between hand and foot, but the implementation of such mechanism is limited by practical factors. These possibilities are not mutually exclusive, and can be addressed by future studies by either testing the effect of training on foot grasping, or testing special populations who are experienced with foot actions (e.g. people born without hands; Striem-Amit, Vannuscorps, & Caramazza, 2017, Dempsey-Jones et al., 2019, Liu et al., 2020).

Finally, we note the possibility that difference in the scaling of aperture size to object size between hand and foot may reflect distinct visuomotor representations. Past studies provide evidence for distinct visuomotor representations between typical grasping with the thumb and index finger, and novel grasping with the thumb and little finger (Gonzalez et al., 2008). Therefore, it is possible that perceptual information about object size may not be similarly or efficiently encoded by the visuomotor system during early phase of foot actions (Freud et al., 2016), generating less coordinated reaching and preshaping components. Our data does not allow fully distinguish what visuomotor representations underlie foot and hand actions respectively, and future studies can investigate this question by studying how perceptual information is represented during hand and foot actions, e.g., by using neuroimaging methods.

### Summary

We tested whether the characteristic kinematic profile of hand reaching and grasping may be driven by motor control unique to the hands, or if similar kinematics can be found for the foot, an untrained and remote body part. We found similar velocity profile during the acceleration phase between the hand and foot, likely reflecting common motor plan mechanisms. In addition, maximum grip aperture scaled with object size for both hand and foot, indicating a common grasping control mechanism that takes into account object size during pre-shaping. One remarkable difference between hand and foot lies in longer deceleration phase for foot that may reflect overall higher accuracy demand on the foot. Additionally, sensitivity to object size was less elaborate and manifested much later for foot vs. hand. These findings also point to different temporal coupling of transport and manipulation between foot and hand. It is not clear from our data whether these differences reflect distinct neural mechanisms, biomechanical constraints in the foot, or lack of experience in foot motor control of participants. Although future studies are required to address the role of experience on some parameters of temporal features of foot grasping, our data provides support for shared kinematics and motor control planning between the hand and the foot. Together with recent imaging studies, this supports the existence of some effector-independent neural representations of action, that do not rely on shared use and experience.

