## Supplementary figure S1 for "Are reaching and grasping effector-independent? Similarities and differences in reaching and grasping kinematics between the hand and foot"

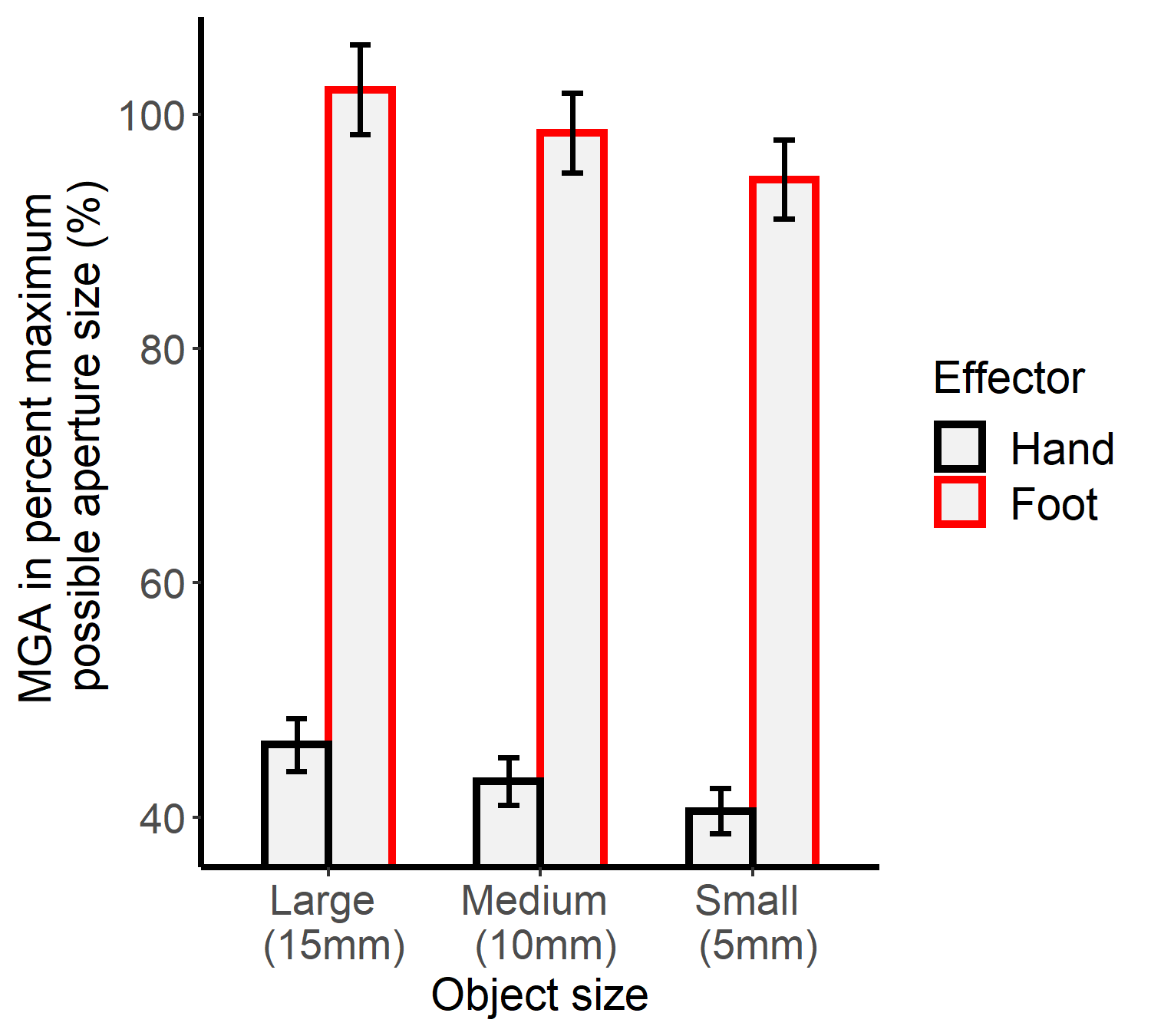

Figure S1. Maximum aperture size (MGA) during grasping expressed as percentage maximum possible aperture size in each effector. Whereas the MGA of the hand is far from the limit, the foot MGA is reaching the ceiling of the possible aperture size for all object sizes, indicating a greater difficulty for the foot to perform the grasping task. Error bars denote standard errors.
